# Soybean Cyst Nematode-Resistant Protein AAT_Rhg1_ Affects Amino Acid Homeostasis and Betalain Accumulation

**DOI:** 10.1101/2024.11.14.623632

**Authors:** Yulin Du, Soyoung Jung, Hiroshi Maeda, Andrew F. Bent

## Abstract

Amino acid transporters play crucial roles in plant nitrogen metabolism but also defense responses. AAT_Rhg1_, an apparent amino acid transporter encoded by *Glyma*.*18g022400* (*Rhg1-GmAAT*) at the soybean *Rhg1* locus, contributes to resistance to soybean cyst nematode (SCN), although the *in planta* function of AAT_Rhg1_ remains elusive. In this study we discovered that overexpression of *Rhg1-GmAAT* in soybean roots enhances the betalain pigment synthesis driven by a *RUBY* transgene cassette, potentially through its transporter activity affecting tyrosine level and amino acid homeostasis. Silencing *Rhg1-GmAAT* also moderately increased betalain accumulation while co-overexpression of *Rhg1-GmAAT* and *GmRBOHG* (encoding an AAT_Rhg1_-interacting NADPH oxidase) blocked the betalain phenotype, indicating a complex role of AAT_Rhg1_ in regulating cellular metabolism. Soybean AAT_Rhg1_ did not show a betalain accumulation phenotype when co-overexpressed with *RUBY* in *Nicotiana benthamiana* leaves, suggesting that soybean AAT_Rhg1_ functions differently in *N. benthamiana*. In soybean, the expression of AAT_Rhg1_ proteins mutated at conserved residues D122A or Y268L mitigated or enhanced the betalain phenotypes, respectively, suggesting that these residues are important for AAT_Rhg1_ function. This study advances our understanding of AAT_Rhg1_ while presenting a novel strategy for enhancing betalain biosynthesis by modulating transport and homeostasis of amino acids.

## Introduction

Amino acid transporters play fundamental roles in nitrogen uptake, exchange and distribution in plants (Ortiz-Lopez, 2000). These transporters are essential for plant growth, development, reproduction, and responses to both biotic and abiotic stresses (Yao et al., 2020). In plant-microbial interactions, amino acid transporters and amino acid homeostasis can impact plant defense signaling pathways. During infection, the expression of amino acid transporters in plants can be modulated either as a consequence of pathogenesis or as part of the plant’s defense mechanism (Sonawala et al., 2018). For example, members of the Arabidopsis *Amino Acid Permease* (*AAP*) gene family, which are dynamically regulated upon pathogen infestation, have been associated with susceptibility to infections by root knot nematode (*Meloidogyne incognita*) (Marella et al., 2013) or sugar beet cyst nematode (*Heterodera schachtii*) (Elashry et al., 2013). Additionally, knocking out the Arabidopsis *Lysine Histidine Transporter 1* (*LHT1*) enhances salicylic acid (SA)-dependent broad-spectrum resistance against various bacterial and fungal pathogens (Liu et al., 2010). Glutamine homeostasis, which is influenced by LHT1 has been shown to affect cellular redox status and impact plant disease responses (Liu et al., 2010). It has also been reported that the perception of microbe-associated molecular patterns (MAMPs) in Arabidopsis triggers free amino acid accumulation in an SA-dependent manner, which increases leaf apoplastic amino acids suppressing bacterial virulence and contributes to disease defense responses (Zhang et al., 2023).

AAT_Rhg1_ (encoded by *Glyma*.*18G022400 or “Rhg1-GmAAT”*) is a putative amino acid transporter encoded at the complex *Rhg1* (Resistance to *Heterodera glycines* 1) locus in soybean (*Glycine max*) (Bent, 2022). The *Rhg1* locus carries three tightly linked genes that each contribute to resistance against soybean cyst nematode (SCN, *Heterodera glycines*) (Cook et al., 2012). Copy number variation of the ∼30kb multi-gene *Rhg1* segment significantly impacts SCN resistance (Cook et al., 2012, 2014; Lee et al., 2015; Yu et al., 2016; Patil et al., 2019; Huang et al., 2021). The soybean variety Williams 82 (Wm82) and most other older soybean varieties carry a single copy *Rhg1* segment and are susceptible to SCN, whereas SCN-resistant varieties that carry the *rhg1-b* haplotype (derived from soybean PI 88788) carry as many as ten direct repeat copies of *Rhg1* (Cook et al., 2012, 2014; Lee et al., 2015; Patil et al., 2019). The high-copy *rhg1-b* resistance has been extensively utilized in the field to manage SCN (McCarville et al., 2017). Loss of function experiments have demonstrated that *Rhg1-GmAAT* contributes to resistance against SCN in soybean (Cook et al., 2012). However, despite its importance in genetically encoded SCN resistance, the molecular mechanism underlying SCN resistance and the amino acid transporter activity of AAT_Rhg1_ are not yet fully elucidated.

AAT_Rhg1_ contains eleven transmembrane domains and is predicted to be a member of the tryptophan/tyrosine transporter family (InterPro IPR013059, see also Pfam PF03222) (Paysan-Lafosse et al., 2023). AAT_Rhg1_ has been reported to promote glutamic acid tolerance and impact glutamate transportation in soybean (Guo et al., 2019). However, its mechanistic role in glutamate transport remains unclear. Upon SCN infestation of soybean roots, AAT_Rhg1_ accumulates along the SCN penetration path, with the affected root cells exhibiting an increased abundance of vesicles, large vesicle-like bodies (VLB), and multivesicular and paramural bodies where AAT_Rhg1_ protein is often localized (Han et al., 2023). In *Nicotiana benthamiana*, overexpression of *Rhg1-GmAAT* induces oxidative stress and promotes the formation of large VLB (Han et al., 2023). AAT_Rhg1_ interacts with the soybean NADPH oxidase GmRBOHG (Respiratory Burst Oxidase Homologue G, *Glyma*.*06G162300*) that was previously found to be upregulated upon SCN infection in SCN-resistant soybean (Liu et al., 2019; Han et al., 2023). Co-overexpression of *Rhg1-GmAAT* and *GmRBOHG* in *N. benthamiana* stimulates greater ROS generation (Han et al., 2023). Given that SCN causes the most yield loss of any pathogens of soybean (Bradley et al., 2021), investigating the function of AAT_Rhg1_ protein could provide insights for enhancing SCN resistance in soybean and other potential plant disease resistance mechanisms.

The amino acid transporters that share structural similarities with AAT_Rhg1_ have been identified as essential proteins across various organisms, including yeast, plants, and animals. These transporters play critical roles in maintaining amino acid homeostasis, supporting growth, and mediating responses to environmental stimuli across diverse biological systems. The Arabidopsis ortholog of soybean AAT_Rhg1_ is AtAVT6C (*AT3G56200*), whose function remains largely unexplored. However, research on a closely related protein, AtAVT6D, has shown its involvement in the uptake of aspartic acid in *Xenopus* oocytes (Dhatterwal et al., 2022). Yeast (*Saccharomyces cerevisiae*) AVT6 has been reported to localize to the vacuolar membrane, where it functions as an exporter of aspartate and glutamate from the vacuole (Russnak et al., 2001; Chahomchuen et al., 2009). Human SLC38A5 (solute carrier family 38 member 5) and SLC38A7 (solute carrier family 38 member 7) are orthologous to yeast AVT6. SLC38A5 has been shown to be upregulated and be involved in cell proliferation in both triple-negative breast cancer and pancreatic ductal adenocarcinoma cells (Ramachandran et al., 2021; Sniegowski et al., 2023). SNAT 7 (Sodium-coupled amino acid transporter 7) encoded by *SLC38A7* has been identified as an effluxer of glutamine from lysosomes, which is related to cancer cell growth (Verdon et al., 2017). Another protein from the same family, human SLC38A9, functions as a lysosomal transporter that meditates the efflux from the lysosomal lumen of leucine as well as other non-polar essential amino acids and tyrosine, thereby regulating lysosomal amino acid concentrations (Wyant et al., 2017). Additionally, SLC38A9 senses luminal arginine and activates the mTORC1 (mechanistic target of rapamycin complex 1) to regulate cell growth (Wyant et al., 2017). Due to their roles in pathogenic cellular processes, these human amino acid transporters are considered important potential targets for disease therapy. Given the high-confidence of the predicted structures (Jumper et al., 2021) and close structure alignment (Bittrich et al., 2024) of AAT_Rhg1_ with yeast AVT6, human SLC38A7 (SNAT7) and zebrafish SLC38A9 (PDB 6C08; Lei et al., 2018), which have been demonstrated to possess amino acid transporter activities, we hypothesize that AAT_Rhg1_ also functions as an amino acid transporter (Russnak et al., 2001, Chahomchuen et al., 2009; Hägglund et al., 2011; Rebsamen et al., 2015).

*RUBY* is an artificial gene cassette that drives the production of betalain pigments derived from the aromatic amino acid precursor tyrosine, resulting in a distinct red coloration in plant tissues where it is expressed (He et al., 2020). *RUBY* has been used as a screenable visual marker of transgenic plants and a noninvasive gene expression monitor in various plant species. Betalains, most well-known from beets (*Beta vulgaris*), are subclassified into yellow betaxanthins and red-violet betacyanins, which have strong antioxidant activity and nutritional values (Polturak, 2018). Betalains are widely used as natural food colorants. Due to environmental and potential health concerns regarding synthetic food dyes derived from petroleum, demand for betalains has been increasing (Downham & Collins, 2000; Wrolstad & Culver, 2012). To improve betalain production, research and breeding efforts have focused on promoting betalain biosynthesis in not only red beet, but also in alternative plant species, tissue culture, cell culture lines, microbes and bioreactor systems (Polturak and Aharoni, 2018; Murthy et al., 2024). In the present study we initially used *RUBY* as the screenable marker to track positive transformants in our experiments that co-overexpressed *Rhg1-GmAAT* and *GmRBOHG* for SCN resistance tests. Unexpectedly, we discovered that co-overexpression of *Rhg1-GmAAT* and *RUBY* enhanced betalain accumulation in soybean roots. We hypothesized that the enhanced betalain accumulation is associated with AAT_Rhg1_ amino acid transporter activity and carried out multiple lines of study to dissect this unexpected betalain phenotype and the elusive functions of AAT_Rhg1_.

## Materials and Methods

### Plasmid constructs

To silence *Rhg1-GmAAT*, a 292bp fragment bridging *Rhg1-GmAAT* CDS1 and CDS2 was used to generate the hairpin RNA construct. This fragment and its reverse complement were cloned into the pCambia2300 plasmid driven by *GmUbi*_*pro*_ and an *Rbcs-E9* terminator. A CaMV *35S* promoter driven spectinomycin resistance marker with chloroplast targeting signal was used for transgenic soybean screening (Martinell et al., 2013). In the overexpression constructs, *RUBY, Rhg1-GmAAT*, and *GmRBOHG* were overexpressed utilizing double CaMV *35S* promoter with TMV omega enhancer (pICH51288), and nopaline synthase (NOS) terminator. *Rhg1-GmAAT* and *GmRbohG* coding sequence were derived from constructs used in Han et al. (2023). Plasmids containing multiple overexpression modules were cloned using Golden Gate Assembly into binary vector pAGM4673 (MoClo Tool Kit) (Weber et al., 2011). BbsI cutting sites or Gibson Assembly overhangs were added via PCR to gene modules to be cloned into Golden Gate level 0 plasmids. Since *GmRbohG* is incompatible with Golden Gate Assembly, it was inserted into the final binary vector using customized ApaI and PmeI enzyme sites at position 2 using T4 DNA ligase. The *RUBY* construct was derived from Addgene plasmid #160908. AAT_Rhg1_ single amino acid mutations were made using site directed mutagenesis with overlapping oligonucleotide primers and DpnI digestion of template DNA. A list of the oligonucleotide primers used for RT-qPCR and DNA plasmid construction is provided as Supplemental Table 1.

### Transgenic plant materials

Empty vector (EV) and *Rhg1-GmAAT* RNAi silencing constructs were transformed into soybean variety IL3025N at the UW-Madison Wisconsin Crop Innovation Center (WCIC). Stable homozygous transgenic lines were identified and 4^th^ generation (T4) plants were used. Composite soybean plants with non-transgenic shoots and transgenic roots were generated using the methods described in Estrada-Navarrete & Alvarado-Affantranger (2007) and Fan et al.(2020). In overexpression experiments, composite plant transgenic soybean root samples for RNA extraction were taken 21 days after transformation. Composite plants were grown for another 30 days for betalain extraction. In the silencing experiment, composite plant transgenic root samples were taken for RNA or betalain quantification 40 days after transformation. Detached transgenic soybean roots were generated using the method described in Cook et al. (2012), and samples for RNA and metabolite extractions were taken 35 days after transformation. Transient overexpression in *N. benthamiana* leaf was conducted according to Li (2011). Images of infiltrated leaves were taken 48 hours after infiltration, followed by sample collection from infiltrated leaves for betalain quantification.

### RNA extraction and RT-qPCR

Sampled soybean root tissues were collected and immediately flash-frozen in liquid nitrogen and stored at -80°C before RNA extraction. Frozen samples were homogenized with 2mm zirconium silicate grinding beads and PowerLyzer™ 24 (MO BIO Laboratories, Inc.). Plant RNA was extracted using Direct-zol™ RNA MiniPrep Plus Kits from Zymo Research following the manufacturer’s instructions. cDNA was synthesized using FIREScript^®^ RT cDNA synthesis MIX from Solis BioDyne. quantitative reverse transcription polymerase chain reaction (RT-qPCR) was conducted using HOT FIREPol^®^ EvaGreen^®^ qPCR Supermix from Solis BioDyne and CFX96 real-time PCR detection system (BioRad, Hercules, CA).

### Metabolite extraction and quantification

Sampled plant tissues were immediately flash-frozen in liquid nitrogen and stored at -80°C, except for the betalain experiment with *Rhg1-GmAAT* RNAi and corresponding empty vector soybean lines, for which root samples were freeze-dried.

For soybean root samples, the levels of tyrosine and other amino acids were analyzed as in our prior studies (Jung and Maeda, 2024). Approximately 30 mg of ground frozen samples were rapidly mixed by vortexing with 400 μl of chloroform:methanol (2:1, v/v) metabolite extraction buffer together with 0.5 μg/ml of isovitexin as an internal standard. Then, 300 μl of water and 125 μl of chloroform were added, and samples were vortexed vigorously and centrifuged at 10,000 g for 5 min. Subsequently, 450 μL of the polar phase was transferred to a fresh tube, dried down overnight in a SpeedVac at room temperature, and resuspended into 100 μl of liquid chromatography-mass spectrometry (LC-MS) grade water for LC-MS analysis and total betacyanins level quantification.

LC-MS analysis of plant metabolites was performed using a Vanquish Horizon Binary UHPLC (Thermo Scientific, Waltham, MA, USA) coupled to a Q Exactive Orbitrap mass spectrometer (Thermo Scientific, Waltham, MA, USA), One micro liter of each resuspended sample was injected onto a HSS T3 C18 reversed phase column (100 × 2.1 mm i.d., 1.8 μm particle size; Waters, Milford, USA) and eluted using a 25-minute gradient comprising 0.1 % (v/v) formic acid (Fisher Scientific, Hampton, NH, USA) in LC-MS grade water (solvent A) and 0.1 % (v/v) formic acid in 90% (v/v) LC-MS-grade acetonitrile (solvent B; Fisher Scientific, Hampton, NH, USA) at a flow rate of 0.4 mL/min and column temperature of 40 °C. The binary linear gradient with the following ratios of solvent B was used: 0-1 min, 1 %; 1-10 min, 1-10 %; 10-13 min, 10-30 %; 13-14.5 min, 30-70 %; 14.5-15.5 min, 70-99 %; 15.5-20 min, 99 %; 20-20 min, 99-1 %; 20-25 min, 1 %. MS spectra were recorded using full scan in positive mode, covering a mass range from 100 to 1000 mass/charge ratio (m/z). The resolution was set to 140,000, and the maximum scan time was set to 200 ms. The transfer capillary temperature was set to 150°C, while the heater temperature was adjusted to 300°C. The spray voltage was fixed at 3 kV, with a capillary voltage and a skimmer voltage of 25 and 15 V, respectively. The identity of tyrosine, phenylalanine, methionine, leucine, and isoleucine peaks were confirmed by comparing their accurate masses and retention times with those of the corresponding authentic standards. The quantification was based on the standard curves for each amino acids generated by injecting different concentrations of authentic chemical standards. Since an authentic standard for tryptophan was not available at that time, the peak area matching with the accurate mass of tryptophan was quantified in relative units normalized by mass of fresh tissue.

For quantification of total betacyanin levels in soybean root samples, 30 μL of the same resuspended samples used for the LC-MS analysis were transferred to individual wells of a 96 transparent plate, flat bottom, half well size (#675101, Greiner Bio-One, Kremsmünster, Austria) with five two-fold serial dilutions. The absorbance at 538 nm was measured using a plate reader (Infinite 200 PRO, TECAN), and multiple data points in the linear range were used to quantify total betacyanin contents. These absorbance values were converted to betacyanin contents using the molar extinction coefficient ε = 60,000 M^−1^ cm^−1^. Betalain pigment extraction and quantification from *N*.*benthamiana* leaves were conducted using the method described in Chang et al. (2021). The absorbance at 538nm and 476nm for betacyanins and betaxanthins respectively were measured spectrophotometrically.

## Supplemental materials

**Supplemental Figure S1**. Other AAT_Rhg1_ single amino acid mutants that did not differently affect betalain accumulation levels.

**Supplemental Figure S2**. Correlation between tyrosine levels and other amino acids upon overexpression of AAT_Rhg1_ WT or its mutants.

**Supplemental Table 1**. Primers used for cloning and RT-qPCR

## Results

### Overexpression of *Rhg1-GmAAT* enhanced the accumulation of betalains synthesized by *RUBY* in soybean roots

The *RUBY* cassette was used as a screenable marker when generating composite plants with transgenic soybean roots where *Rhg1-GmAAT* and/or *GmRBOHG* were overexpressed. Because *Rhg1-GmAAT* contributes to SCN resistance, the overexpression (OE) was conducted in both SCN-susceptible Williams 82 (Wm82) and SCN-resistant Fayette soybean varieties, which respectively carry the single-copy *rhg1-c* or ten-copy *rhg1-b* haplotypes. Surprisingly, when co-overexpressing *Rhg1-GmAAT* with *RUBY* from the same plasmid, visible betalain levels were noticeably higher compared to roots transformed with *RUBY*-only empty vector alone (*RUBY* EV; Figure 1A and 1B). The elevated betalain pigmentation observed upon co-overexpression of *Rhg1-GmAAT* with *RUBY* also was more intense than that observed with *RUBY* + OE *GmRBOHG* and with *RUBY* + OE *Rhg1-GmAAT* + OE *GmRBOHG* co-overexpression constructs (Figure 1A and 1B). Similar results were observed in the backgrounds of both Wm82 and Fayette soybean varieties. The overexpression of *Rhg1-GmAAT* was confirmed by RT-qPCR (Figure 1C and 1D). In roots transformed with the four different constructs mentioned above, *RUBY* transcript levels did not show statistically significant differences (Figure 1E and 1F), indicating that the greater betalain accumulation in *RUBY* + OE *Rhg1-GmAAT* roots was not caused by an elevated *RUBY* mRNA abundance. In both Wm82 and Fayette varieties, the betalain accumulation induced by *Rhg1-GmAAT* overexpression was abolished when co-overexpressing with the AAT_Rhg1_ interactor GmRBOHG (Figure 1A and 1B), suggesting that the physical interaction between AAT_Rhg1_ and GmRBOHG (Han et al., 2023) alters the AAT_Rhg1_-induced increase in betalain accumulation without impacting *RUBY* gene expression.

**Figure 1.**
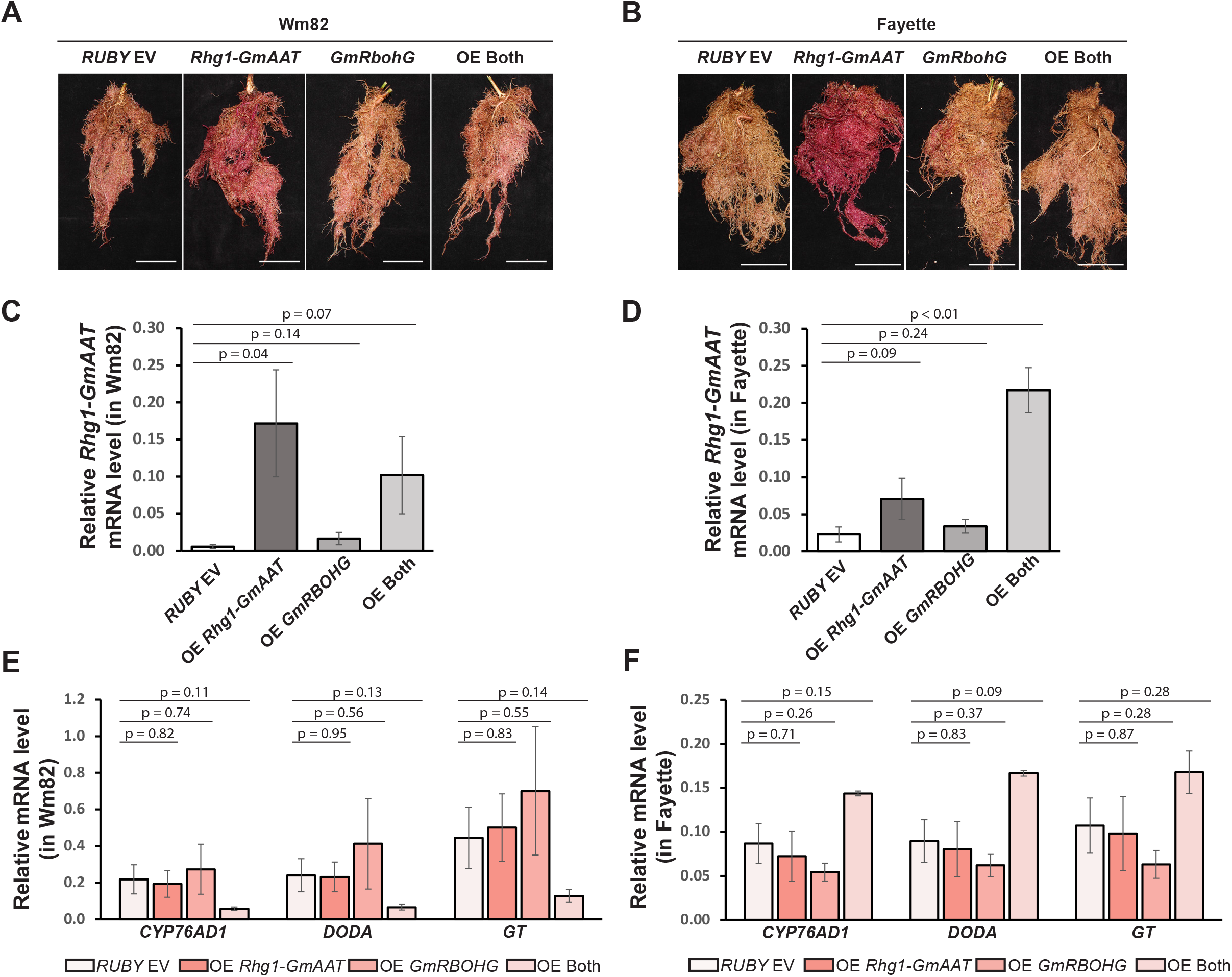
Overexpression of *Rhg1-GmAAT* in soybean roots enhanced the accumulation of betalain synthesized by *RUBY*. **(A, B)** Representative images of transgenic soybean roots from composite plants expressing one of four constructs, including *RUBY* only empty vector *(RUBY* EV*), RUBY* with overexpression of *Rhg1-GmAAT* (*Rhg1-GmAAT*), *RUBY* with overexpression of *GmRBOHG* (*GmRBOHG*), or *RUBY* with overexpression of both *Rhg1-GmAAT* and *GmRBOHG* (OE Both), in soybean variety Williams 82 (Wm82) **(A)** or Fayette **(B)**. Scale bars = 5cm. **(C, D)** Transcript levels of *Rhg1-GmAAT* relative to *GmEF1A* in Wm82 **(C)** or Fayette **(D)** transgenic roots expressing the same four constructs respectively as described above. Data were analyzed using one-tailed *t*-test (n = 3) to compare to *RUBY* EV and presented as mean ± SE. **(E, F)** Transcript levels of three *RUBY* genes (*CYP76AD1, DODA* and *GT*) relative to *GmEF1A* in Wm82 **(E)** or Fayette **(F)** transgenic roots expressing the four constructs respectively. Data were analyzed using two-tailed *t*-test (n = 3) to compare to *RUBY* EV and presented as mean ± SE.

We expected that AAT_Rhg1_ would also enhance betalain accumulation when *Rhg1-GmAAT* was expressed with *RUBY* in *N. benthamiana* leaves. However, in *N. benthamiana*, transient overexpression of the *RUBY*-containing constructs showed phenotypes opposite to those observed in soybean roots. The *RUBY*-only construct resulted in the highest betalain accumulation (Figure 2A and 2B), while the co-overexpression constructs exhibited significantly lower betalain levels. Although the difference was not statistically significant, *RUBY* + OE *Rhg1-GmAAT* produced even lower betalain level compared to *RUBY* + OE *GmRBOHG*, and to the *RUBY* + OE *Rhg1-GmAAT* + OE *GmRBOHG* co-overexpression construct, as well as to *RUBY* + OE *Rhg1-GmAAT* and *RUBY* + OE *GmRBOHG* co-infiltration. This suggests that soybean AAT_Rhg1_ functions differently in *N. benthamiana* leaves.

**Figure 2.**
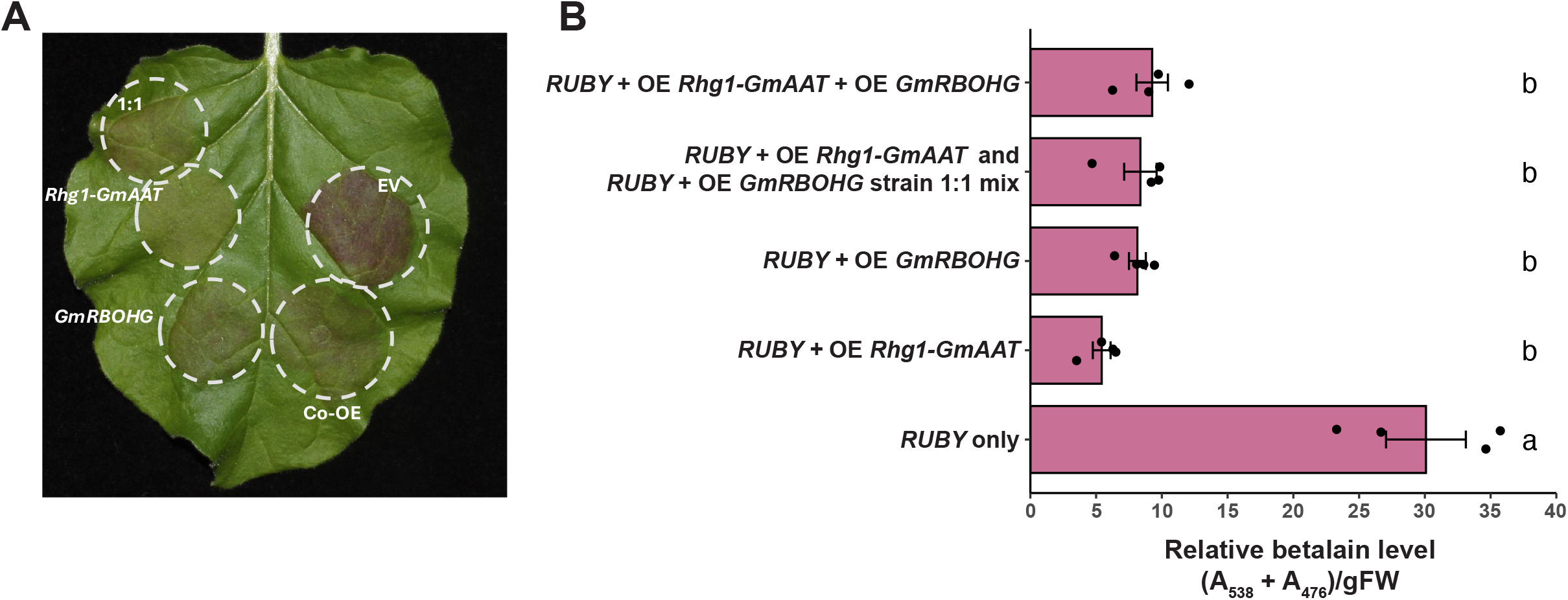
Overexpression of *Rhg1-GmAAT* in *Nicotiana benthamiana* leaves reduced the accumulation of betalain synthesized by *RUBY*. **(A)** Representative image of *N. benthamiana* leaves infiltrated with Agrobacterium carrying various *RUBY* constructs: a 1:1 mixture of *RUBY* + OE *Rhg1-GmAAT* and *RUBY* + OE *GmRBOHG* Agrobacterium strains (labeled as 1:1), *RUBY* + OE *Rhg1-GmAAT* alone, *RUBY* + OE *GmRBOHG* alone, *RUBY +* OE *Rhg1-GmAAT* + OE *GmRBOHG*, and *RUBY* only (empty vector, EV). **(B)** Quantification of relative betalain levels as (A_538_ + A_476_) per gram of fresh weight (gFW) in infiltrated leaf tissues. Data were analyzed using ANOVA and Tukey HSD (n = 4), and are shown as mean ± SE.

### Silencing *Rhg1-GmAAT* in soybean also moderately increased betalain accumulation

To further examine the effects of *Rhg1-GmAAT* expression on betalain pigment accumulation in soybean roots, *RUBY* was expressed in composite soybean plants. Transgenic roots expressing *RUBY* were grown from soybean plants of *rhg1-b/rhg1-b* genotype already carrying stable transgenic *Rhg1-GmAAT* RNAi silencing or empty vector (EV) constructs. Silencing of *Rhg1-GmAAT* was verified in the stable transgenic lines by RT-qPCR (Figure 3A). Unexpectedly, betacyanins accumulated 2.7-fold more when *Rhg1-GmAAT* was silenced compared to the empty vector line (Figure 3B), mimicking the results from *Rhg1-GmAAT* overexpression (Figure 1A, 1B and 4D). RT-qPCR confirmed that, similar to *Rhg1-GmAAT*-overexpressing roots, this higher betalain level observed in *Rhg1-GmAAT*-RNAi plants was not attributable to elevated *RUBY* mRNA abundance (Figure 3C).

**Figure 3.**
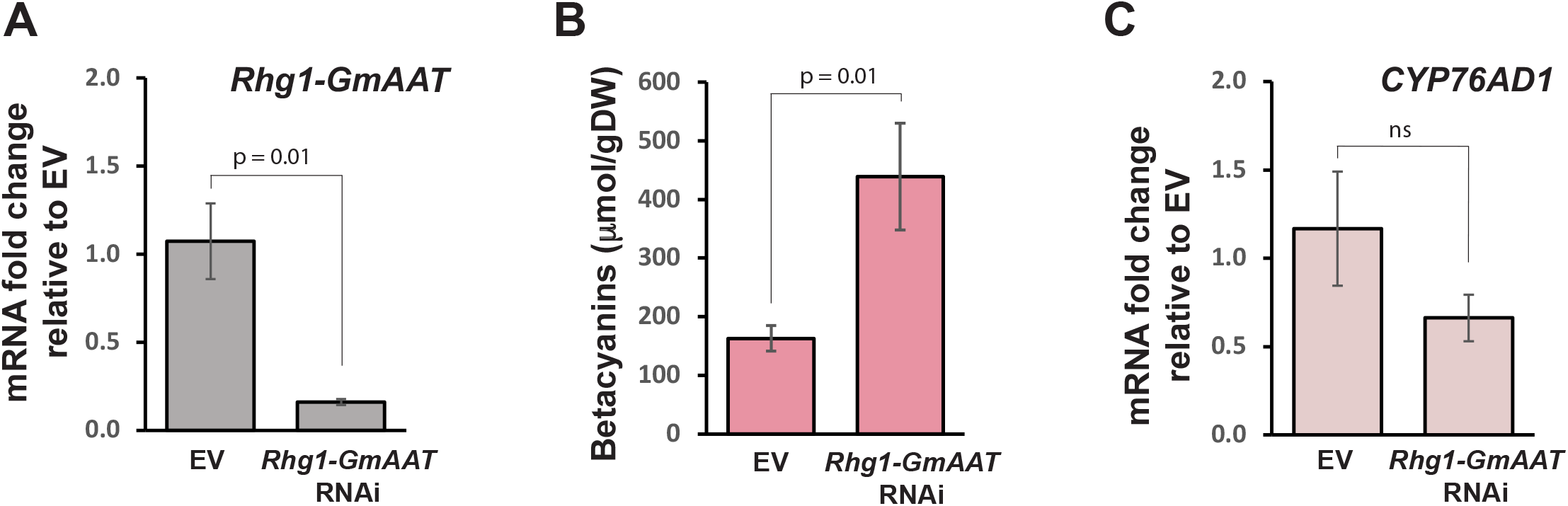
Silencing *Rhg1-GmAAT* in soybean roots moderately enhanced the accumulation of betalains synthesized by *RUBY*. **(A)** Transcript fold change of *Rhg1-GmAAT* relative to mean for empty vector (EV) soybean line, in transgenic EV or *Rhg1-GmAAT*-silencing (*Rhg1-GmAAT* RNAi) soybean roots expressing *RUBY*. Stable transgenic soybean lines were generated by transforming the IL3025N soybean variety with either an EV or *Rhg1-GmAAT* RNAi construct. EV or *Rhg1-GmAAT* RNAi stable transgenic lines were subsequently transformed with the *RUBY* cassette. Data were analyzed using one-tailed *t*-test (n = 5) and shown as mean ± SE. **(B)** Quantification of betacyanin levels per gram of dry weight (gDW) in roots described above. Data were analyzed using one-tailed *t*-test (n > 12) and are shown as mean ± SE. **(C)** Transcript fold change of one of the *RUBY* genes *CYP76AD1* relative to EV soybean line in roots described above. Data were analyzed using two-tailed *t*-test (n = 5) and shown as mean ± SE.

### Y268L or D122A mutations in AAT_Rhg1_ respectively enhance or attenuate *RUBY*-mediated betalain accumulation

To further investigate the function of AAT_Rhg1_ protein in *RUBY*-mediated betalain biosynthesis in soybean roots, we mutated AAT_Rhg1_ residues that are conserved among amino acid transporters and expressed these mutated AAT_Rhg1_ in soybean roots. AAT_Rhg1_ was first aligned with the other 14 annotated amino acid transporters in the soybean Wm82 reference genome (*Glycine max*. Wm82.a6.v1) and with 387 additional annotated amino acid transporters from Viridiplantae gene family 123428317 obtained from associated PlantFAMs via hmmsearch on Phytozome. Based on HMM logos of the alignment results produced by Skylign (Wheeler et al., 2014), four conserved and potential structurally or functionally essential residues (Asp 122, Tyr 268, Asp 286, and Trp 370) were identified and selected for site directed mutagenesis (Figure 4A, 4B and S1A). In addition to substitutions of those conserved residues with small neutral alanine, or large neutral phenylalanine, Tyr 268 was also mutated to leucine and Trp 370 was also mutated to isoleucine. These AAT_Rhg1_ mutants were co-overexpressed with *RUBY* in transgenic roots generated on SCN-susceptible Wm82 soybean composite plants.

**Figure 4.**
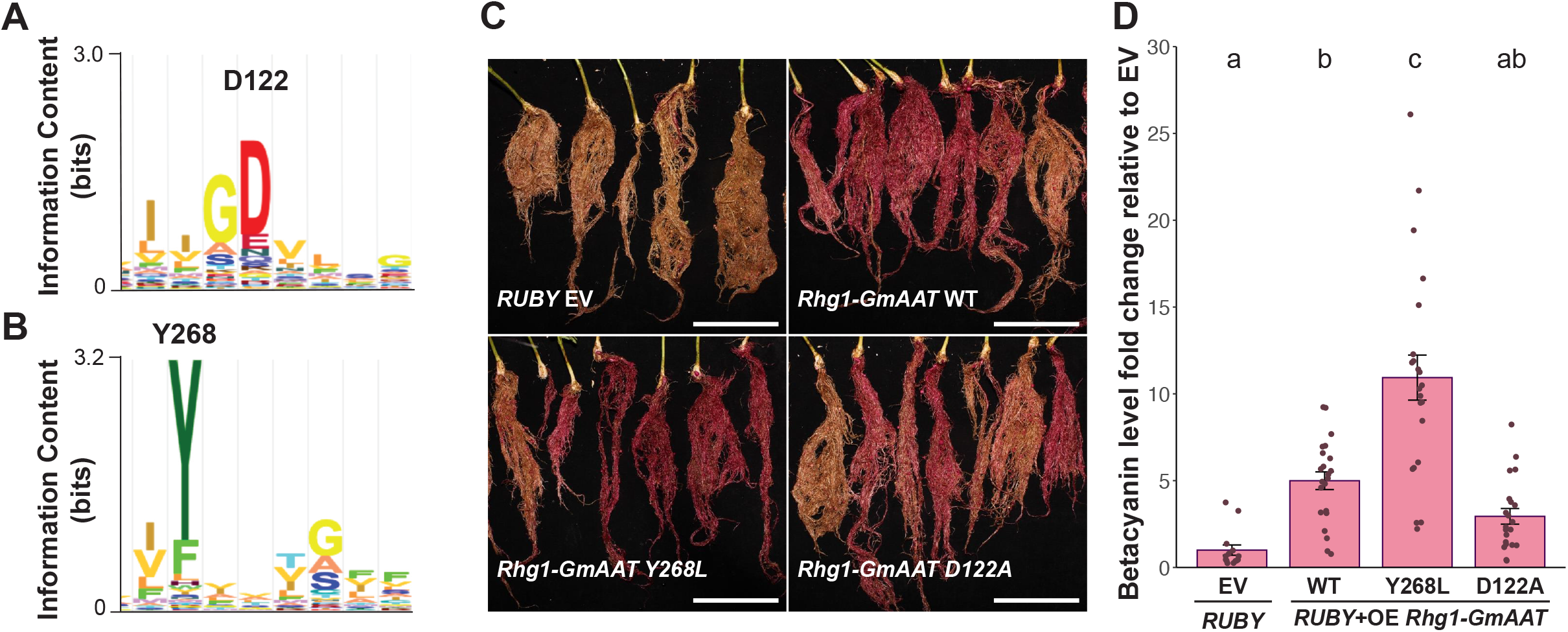
Two AAT_Rhg1_ single amino acid mutants showed altered betalain accumulation levels. **(A, B)** HMM logo showing a portion of the alignment of AAT_Rhg1_ with 387 other amino acid transporters from the Viridiplantae gene family 123428317. Two conserved residues of AAT_Rhg1_, Asp 122 (D122) (A) and Tyr 268 (Y268) (B), were selected for site-directed mutagenesis. **(C)** Representative images of transgenic soybean roots expressing *RUBY* alone (EV), *RUBY* with overexpression of wild-type *Rhg1-GmAAT* (*Rhg1-GmAAT* WT), *RUBY* with overexpression of *Rhg1-GmAAT Y268L*, and *RUBY* with overexpression of *Rhg1-GmAAT D122A* constructs. Scale bars = 10 cm. **(D)** Quantification of betacyanin levels in transgenic roots described in (C). Data from two biological replicates were combined and plotted. A two-way ANOVA showed no significant differences between replicates. The means for bars sharing the same letter were not significantly different (ANOVA Tukey HSD; n > 14). Data are presented as mean ± SE.

As in previous experiments (Figure 1), co-expression of *RUBY* and AAT_Rhg1_ WT (wild type) showed a strong increase in betalain accumulation as compared to the *RUBY*-only control (Figure 4C and 4D). However, the expression of two AAT_Rhg1_ mutants Y268L and D122A impacted betalain levels differently than the non-mutagenized AAT_Rhg1_ WT (Figure 4C and 4D). Compared to AAT_Rhg1_ WT, AAT_Rhg1_ Y268L displayed even stronger betalain accumulation, while AAT_Rhg1_ D122A showed an attenuated betalain level (Figure 4C and 4D). The other mutations at conserved residues, D122F, D286A, D286F, W370A, W370I, Y268A and Y268F, did not cause statistically significant differences relative to non-mutagenized AAT_Rhg1_ (Figure S1B). These results reveal that the Y268L and D122A mutations alter the functionality of AAT_Rhg1_ in opposite directions.

### Overexpression of *Rhg1-GmAAT* affects amino acid homeostasis in soybean roots

Tyrosine is the direct substrate for betalain biosynthesis by *RUBY* (He et al., 2020; Deng et al., 2023). As AAT_Rhg1_ belongs to the tryptophan/tyrosine permease family, we hypothesized that AAT_Rhg1_ affects betalain synthesis by altering the availability of the betalain precursor tyrosine in the cytosol, where betalain biosynthesis takes place (Chen et al., 2017). To test how overexpression of *Rhg1-GmAAT* affects aromatic amino acid levels, metabolites were extracted from detached transgenic soybean roots expressing *Rhg1-GmAAT* WT or the two mutants, *Rhg1-GmAAT D122A* and *Rhg1-GmAAT Y268L*. In this experiment, the constructs were expressed without the *RUBY* cassette, which would further metabolize tyrosine, in order to assess the impacts of the AAT_Rhg1_ variants on amino acid levels without being complicated by their metabolism. The levels of tyrosine, phenylalanine, and tryptophan were quantified from these samples by LC-MS (Figure 5A, 5B, 5C). Phenylalanine levels were significantly lower in roots overexpressing *Rhg1-GmAAT* WT or *Rhg1-GmAAT Y268L* but there was no significant difference between EV and *Rhg1-GmAAT D122A* (Figure 5B). Tyrosine and tryptophan levels did not show statistically significant differences in soybean roots expressing *Rhg1-GmAAT* WT or the mutants, although there was a trend that the overexpression of *Rhg1-GmAAT* WT or *Rhg1-GmAAT Y268L* may have lowered tyrosine and tryptophan abundance (Figure 5A and 5C). *Rhg1-GmAAT Y268L* overexpression (which caused even greater betalain accumulation than *Rhg1-GmAAT* WT when co-expressed with *RUBY*; Figure 5D) did not show a different tyrosine abundance compared to *Rhg1-GmAAT* WT samples (Figure 5A).

**Figure 5.**
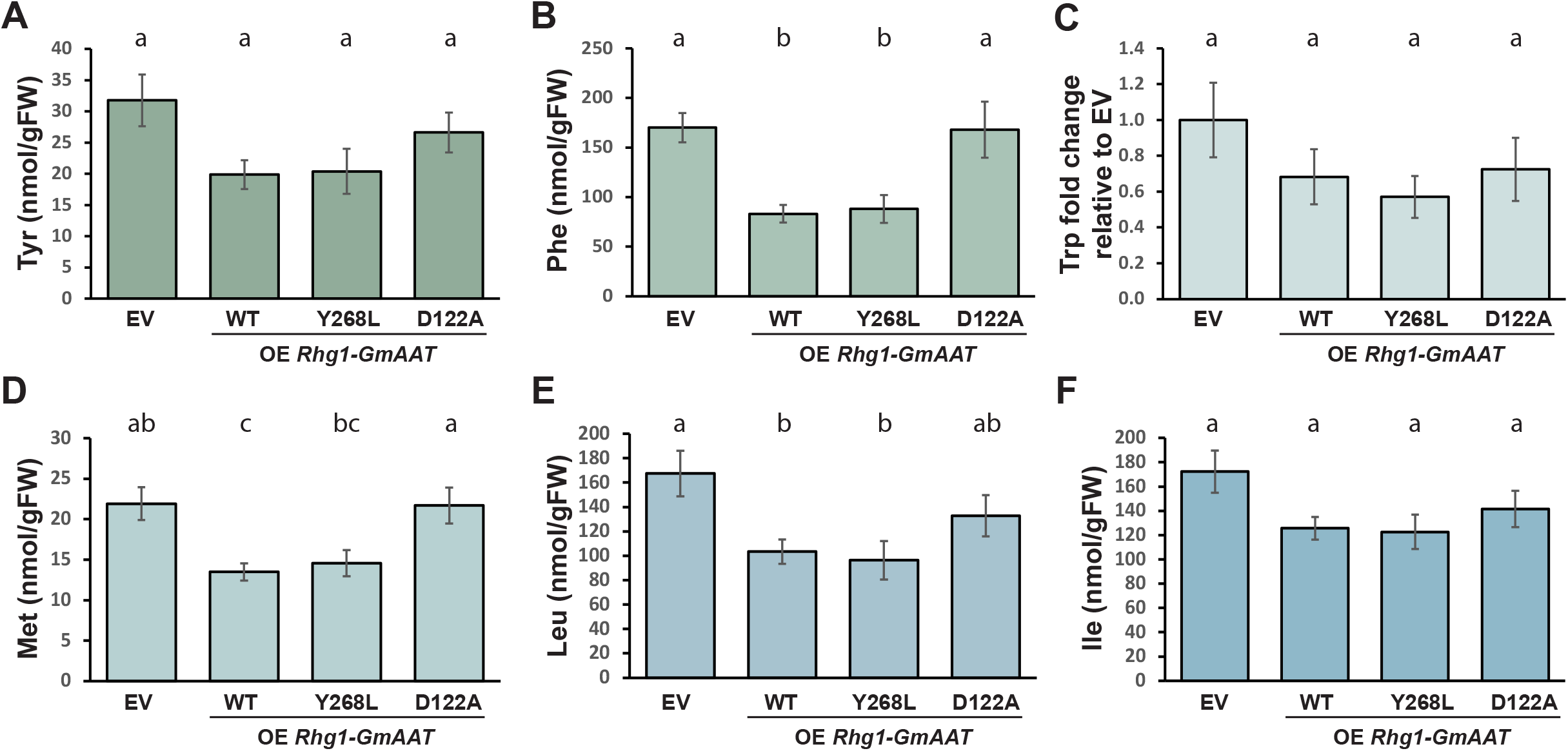
Overexpression of *Rhg1-GmAAT* affected amino acid homeostasis in soybean. Quantification of amino acid levels in detached transgenic soybean roots overexpressing empty vector (EV), *Rhg1-GmAAT* WT, *Rhg1-GmAAT Y268L*, or *Rhg1-GmAAT D122A* constructs, identified using TdTomato as the transgenic screenable marker. The amino acids measured include Tyrosine (Tyr) **(A)**, Phenylalanine (Phe) **(B)**, Tryptophan (Trp) fold change relative to EV **(C)**, Methionine (Met) **(D)**, Leucine (Leu) **(E)**, and Isoleucine (Ile) **(F)**. The means for bars sharing the same letter within a graph were not significantly different (ANOVA Tukey HSD; n > 9). Data presented as mean ± SE.

To examine if the above trends extended beyond aromatic amino acids, changes in methionine, leucine, and isoleucine (which could be quantified from the same LC-MS runs) were also examined (Figure 5D, 5E, 5F). The abundance of these amino acids showed similar trends as tyrosine and phenylalanine. WT or Y268L AAT_Rhg1_ expression reduced the abundance of each of the examined amino acids, with some being statistically significantly. For all six examined amino acids, the expression of AAT_Rhg1_ D122A led to amino acid levels that were never significantly different from the empty-vector negative control, suggesting that this mutated form of AAT_Rhg1_ may be non-functional.

In roots overexpressing *Rhg1-GmAAT Y268L*, tyrosine levels showed a strong positive correlation with all other quantified amino acids (Figure S2). For EV, *Rhg1-GmAAT* WT and *Rhg1-GmAAT D122A*, the correlation with tyrosine levels was strong only for some but not all of the other measured amino acids (Figure S2). Overall, the results suggest that elevated expression of functional forms of *Rhg1-GmAAT* impacted the homeostasis of aromatic amino acids as well as other amino acids.

## Discussion

In this study we demonstrated that the abundance of AAT_Rhg1_, a protein that contributes to SCN resistance, can affect the amount of betalain pigments synthesized by the three *RUBY* enzymes and influence amino acid homeostasis in soybean roots. The results from the tyrosine quantification did not support our hypothesis that the overexpression of *Rhg1-GmAAT* specifically increases total cellular tyrosine levels and thereby enhances betalain synthesis. In plants, tyrosine is synthesized within the plastids from the final product of the shikimate pathway, chorismate. Two key enzymes in this tyrosine biosynthetic pathway, chorismate mutase (CM) and TyrA arogenate dehydrogenase (TyrA_a_), are strongly inhibited by tyrosine. (Eberhard et al., 1996; Rippert & Matringe, 2002; Schenck et al., 2015; Schenck & Maeda, 2018). This negative feedback mechanism is essential for maintaining cellular tyrosine levels and balancing of aromatic amino acids. Based on our findings, we now speculate for future work that the expression of *Rhg1-GmAAT* directly or indirectly increases tyrosine levels in the cytosol.

As metabolite quantification in different subcellular compartments remains challenging, our experiment quantified amino acids from whole-cell extracts rather than certain cellular fractions such as cytosolic or vacuolar extracts. Therefore, a globally lower tyrosine level in OE *Rhg1-GmAAT* soybean root cells does not necessarily indicate reduced tyrosine in the cytosol, where betalain biosynthesis takes place (Chen et al., 2017). CYP76AD1 catalyzes the first step of betalain biosynthesis. Given that CYP76AD1 was shown to localize in the cytoplasm and nucleus in *N. benthamiana* (Chen et al., 2017), betalain levels could serve as an indicator of tyrosine availability in the cytosol. Surprisingly, silencing *Rhg1-GmAAT* also led to increased betalain accumulation. This again might be attributable to the impact of AAT_Rhg1_ on subcellular tyrosine balance and feedback regulation of tyrosine synthesis. According to DeepLoc2.1 tool (Ødum et al., 2024), the tonoplast is predicted to be the most likely subcellular location of AAT_Rhg1_. This suggests that AAT_Rhg1_ may play a role in exporting tyrosine from vacuolar storage and potentially increase cytosolic tyrosine availability for betalain synthesis when *RUBY* is expressed.

Although the exact molecular mechanisms of how AAT_Rhg1_ affects betalain accumulation have not been elucidated, this research offers a possible new strategy to promote the biosynthesis of betalain and other tyrosine-derived compounds, such as antioxidants and defense metabolites. Jung and Maeda (2024) employed a “push-and-pull” strategy to enhance betalain production, by co-expressing a feedback-insensitive arogenate dehydrogenase BvTyrAα and the rate-limiting enzyme DODA in *N. benthamiana*, boosting both tyrosine production and utilization. The present study documented that overexpression of AAT_Rhg1_ can elevate betalain accumulation in its normal host species. We speculate that the observed elevated betalain levels are derived from decreased vacuolar tyrosine storage and accordingly increased availability of cytosolic tyrosine levels. This could be due to the direct transport function of AAT_Rhg1_ or a mechanism that may also be achievable by manipulating other tonoplast-localized amino acid transporters.

We identified two AAT_Rhg1_ single amino acid mutants, D122A and Y268L, that altered the betalain hyperaccumulation phenotype caused by co-overexpression of *Rhg1-GmAAT* and *RUBY*. These two residues are often conserved among amino acid transporters across 388 plant species, suggesting their probable importance in maintaining proper transporter activity. Although the structure of AAT_Rhg1_ remains to be determined, there are reports on crystal structures of similar proteins, including SLC38A9 (Lei et al., 2018), its bacterial homologue LeuT (Krishnamurthy & Gouaux, 2012), the prokaryotic SLC7 homologue GkApcT (Jungnickel et al., 2018), and bacterial BasC (Errasti-Murugarren et al., 2019). Based on the protein structure predicted by AlphaFold2 (Jumper et al., 2021), Asp 122 of AAT_Rhg1_ is located near a hinge loop (G126 – S149) that connects TM3 (transmembrane helix 3) and TM4 and potentially forms a flexible cap over the main channel of AAT_Rhg1_ (Figure S1C). A D122A mutation may alter placement of this hinge structure, potentially blocking the transporter channel or impeding the necessary conformational changes for its function, which would account for the failure of AAT_Rhg1_ D122A to impact the levels of betalains or the measured amino acids (Figure 4 and 5). The space-occupying D122F mutation did not cause the attenuation of betalain levels observed with the D122A mutant. The D122A mutation may also impact the physical interaction between AAT_Rhg1_ and other proteins, potentially influencing amino acid transport as well as betalain production or degradation.

Tyr 268 is highly conserved among plant amino acid transporters and is located deep within the postulated structure of AAT_Rhg1_ on transmembrane helix 7 (TM7). It faces the interior of the transporter channel and is predicted to protrude toward the groove between TM1a and TM1b (Figure S1C), where it may serve as a substrate-binding site. The bacterial BasC Tyr 236 in TM7 sits in a position analogous to AAT_Rhg1_ Tyr 268, and is also conserved in human L-amino acid transporters (LATs). The BasC Y236F mutant and its homologous Y280F mutant in the human Asc-1 transporter both exhibited increased alanine uptake (Errasti-Murugarren et al., 2019). Studies suggested that Tyr 236 affects the apparent substrate affinity asymmetrically across the membrane, and this is likely due to altered conformational transitions in the transport cycle, rather than through changes in substrate binding affinity (Errasti-Murugarren et al., 2019). Notably, expression of the AAT_Rhg1_ Y268L mutant led to further increase in betalain levels that we interpret as a gain-of-function. AAT_Rhg1_ Y268L may alter substrate binding affinity or impact the transporter conformational changes, thereby influencing transporter activity. It is also of interest that AAT_Rhg1_ with mutations in the conserved Asp 286 and Trp 370 residues resembled non-mutagenized AAT_Rhg1_, causing similar elevation of betalain levels, suggesting that, besides its transport activity, AAT_Rhg1_ may indirectly affect functions of other tonoplast transporters, such as through physical association or hetero-complex formation. We are currently conducting experiments to determine whether the D122A and Y268L AAT_Rhg1_ mutants also alter SCN resistance.

This could provide evidence as to whether AAT_Rhg1_-conferred SCN resistance depends on its amino acid transporter activity. If so, there may be opportunities to enhance SCN resistance by modulating AAT_Rhg1_ activity through mutational screening, with the synthetic betalain phenotype serving as a convenient preliminary indicator that is far easier to assay than SCN resistance phenotyping.

Our previous research has shown that AAT_Rhg1_ expression is upregulated upon SCN infestation, with the AAT_Rhg1_ protein accumulating along SCN penetration path (Han et al., 2023). Here we demonstrate that overexpression of *Rhg1-GmAAT* in soybean roots has effects on the abundance of tyrosine but also other amino acids. Impacts of *Rhg1-GmAAT* overexpression on amino acid homeostasis also were reported in a separate study (Guo et al., 2019). In that study, *Rhg1-GmAAT* overexpression promoted soybean plant tolerance to toxic levels of exogenous Glu, but not Gln, Asp and Gly, which was also associated with increased expression of a soybean glutamate receptor-like gene (*Glyma*.*06G233600*) in comparison to non-transgenic wild-type plants. Additionally, soybean plants overexpressing *Rhg1-GmAAT* showed altered free amino acid profiles in leaf and root tissues compared to wild-type plant. Similar to our results, the overexpression of *Rhg1-GmAAT* led to lower levels of Tyr, Phe, Trp, Met, Leu, and Ile in roots (Guo et al., 2019). The role of AAT_Rhg1_ in regulating amino acid homeostasis could provide a potential explanation for the molecular mechanism behind AAT_Rhg1_-conferred SCN resistance. Since amino acid abundance and cellular distribution can affect plant metabolism, this regulation may have direct or indirect impacts on pathogenesis (G. Liu et al., 2010; Sonawala et al., 2018; Zhang et al., 2023)

In summary, we demonstrate an impact of the AAT_Rhg1_ amino acid transporter on betalain biosynthesis, confirm and further dissect impacts on amino acid homeostasis, and identify two key AAT_Rhg1_ residue mutations with opposing phenotypes. This research not only furthers our understanding of the protein AAT_Rhg1_ that is essential for SCN resistance, but also introduces a novel strategy for enhancing betalain pigment production. Deployment of amino acid transporters to modulate amino acid levels may improve betalain yields in various plant systems. These insights hold implications for plant breeding and biotechnology, potentially advancing the sustainable production of valuable natural pigments.

## Acknowledgements

We thank Roger Innes lab for sharing the *RUBY* construct. We thank Dr. Adam Bayless for making the *Rhg-GmAAT* RNAi construct, and thank Dr. Ryan Zapotocny, Dr. Shaojie Han and Emma Nelson for screening stable transgenic *Rhg-GmAAT* RNAi soybean lines. This work was supported by grant from the University of Wisconsin Agricultural Experiment Station Hatch Program award WIS03087.

## Author Contributions

Y.D. designed the research, performed research, analyzed data and wrote the paper. S.J. performed research, analyzed data and wrote the paper. H.M. designed the research, analyzed data and wrote the paper. A.F.B designed the research, analyzed data and wrote the paper.

## Conflict of interest statement

The authors declare that they have no conflict of interest related to this manuscript.

## Notes

### Competing Interest Statement

The authors have declared no competing interest.

