## Supplemental materials for "Soybean Cyst Nematode-Resistant Protein AAT_Rhg1_ Affects Amino Acid Homeostasis and Betalain Accumulation"

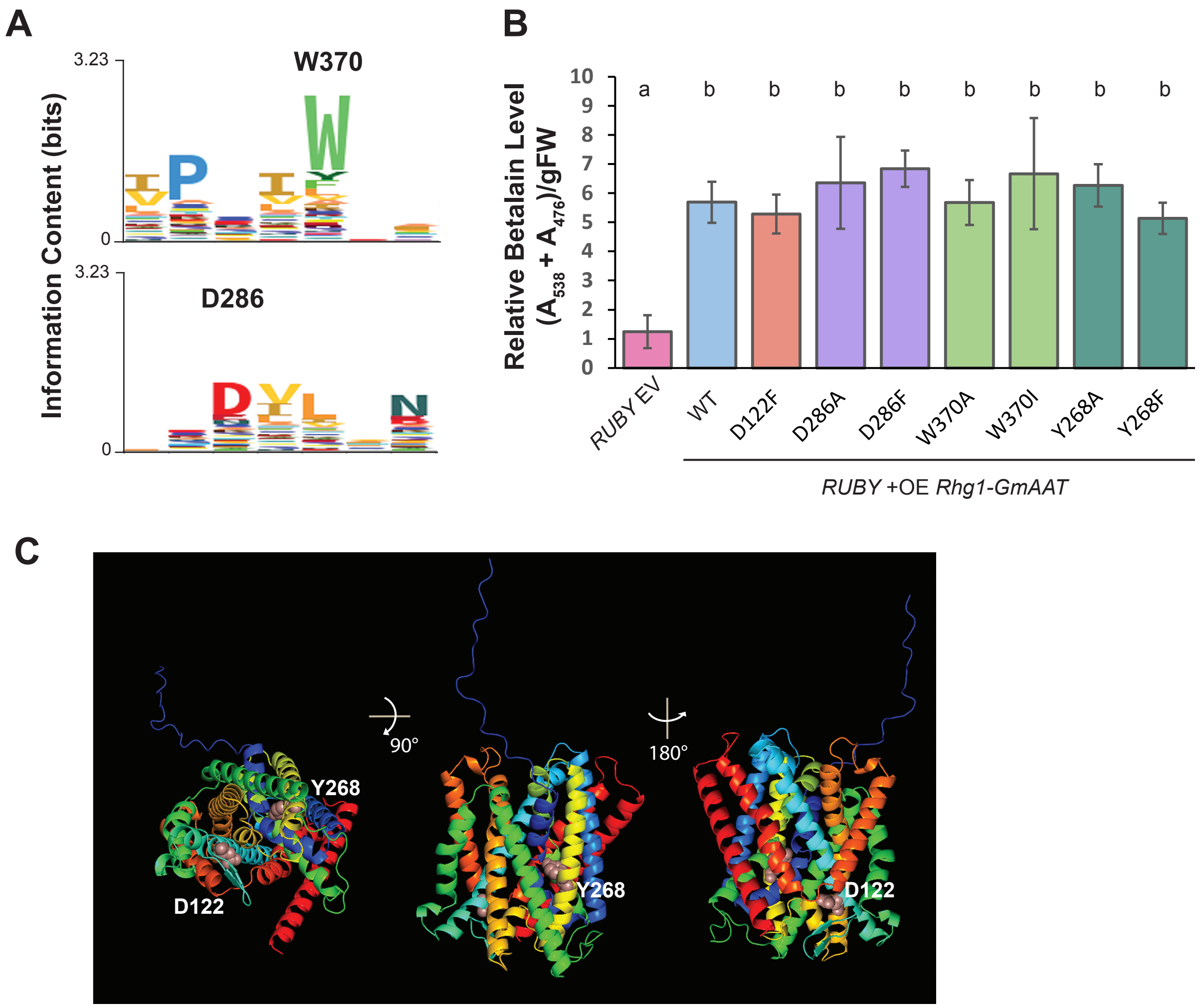

**Supplemental Figure S1. Other $AAT_{Rhg1}$ single amino acid mutants that did not differently affect betalain accumulation levels.**

**(A)** HMM logo showing a portion of the alignment of 15 soybean annotated amino acid transporters. Two conserved residues, Trp 370 (W370) and Asp 286 (D286) were selected for mutagenesis (see also Figure 4).

**(B)** Quantification of betalain levels in transgenic roots expressing *RUBY* empty vector (*RUBY* EV), or *RUBY* with overexpression of *Rhg1-GmAAT* wild type (WT) or mutant constructs. Data were analyzed using ANOVA and Tukey HSD, and are presented as mean ± SE.

**(C)** Predicted structure of the $AAT_{Rhg1}$ protein from AlphaFold2, with the N terminus annotated in blue, C terminus in red, and the Asp 122 (D122) and Tyr 268 (Y268) residues annotated in pink spheres.

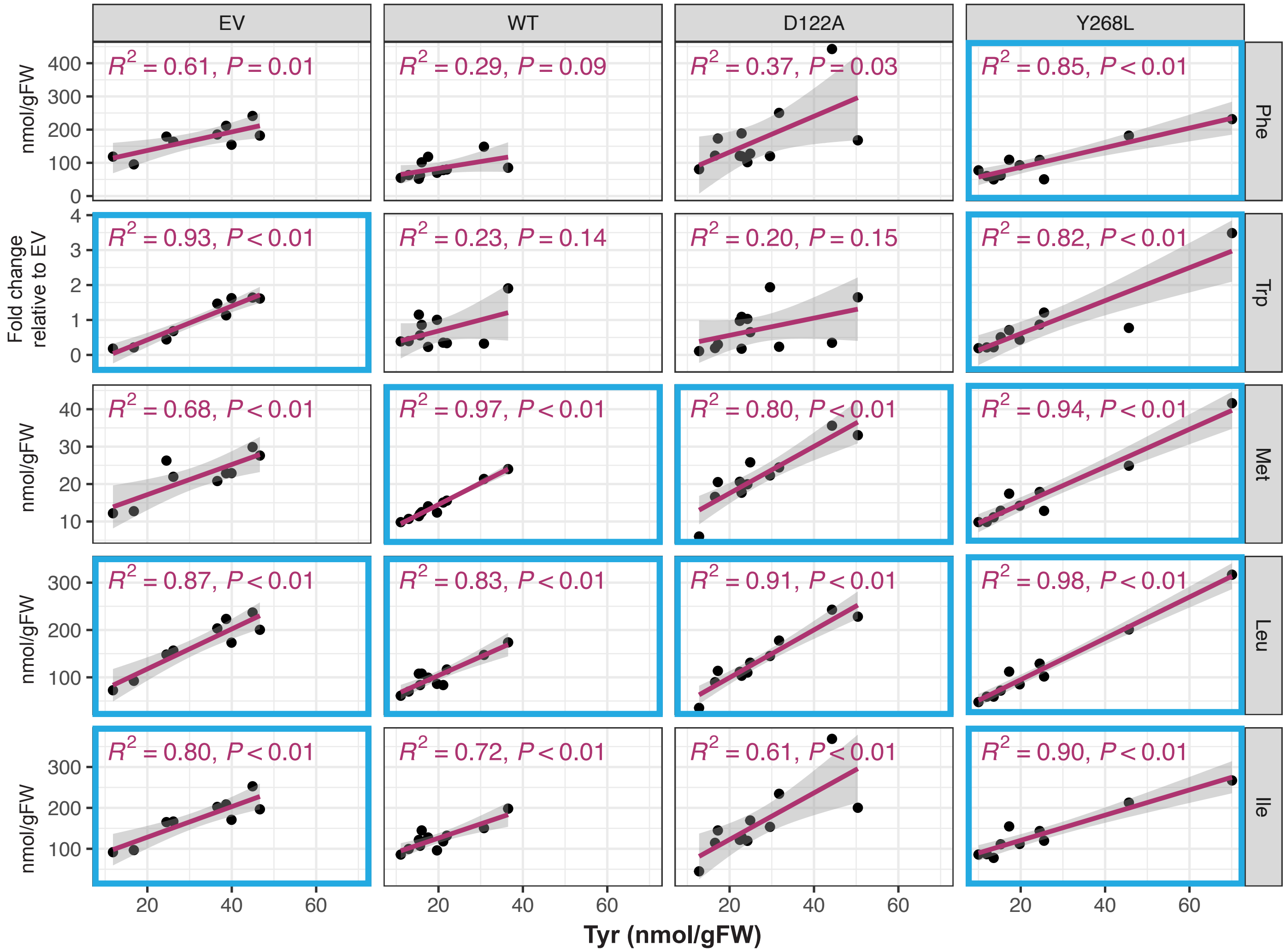

**Supplemental Figure S2. Correlation between tyrosine levels and other amino acids upon overexpression of $AAT_{Rhg1}$ WT or its mutants.**
Linear regression analysis was performed to evaluate the relationship between the abundance of tyrosine (Tyr) and other amino acids, including phenylalanine (Phe), tryptophan (Trp), methionine (Met), leucine (Leu), and isoleucine (Ile), using LC-MS quantification data from transgenic soybean roots overexpressing empty vector (EV), *Rhg1-GmAAT* WT, *Rhg1-GmAAT Y268L*, or *Rhg1-GmAAT D122A* with TdTomato as the transgenic screenable marker. Panels showing strong positive correlations ($R^2 > 0.80$) are highlighted in blue.

**Supplemental Table 1.** Oligonucleotides used for cloning and RT-qPCR.

| | |
|---|---|
| **RT-qPCR primers** | |
| *RUBY_CYP*_qPCR_F | GAGAGACTCGCCCCAGATTCCT |
| *RUBY_CYP*_qPCR_R | CTCGCCCATCGTCAGCTCGTTC |
| *RUBY_DODA*_qPCR_F | TCCGGCCACTGGGAGACAGTGA |
| *RUBY_DODA*_qPCR_R | CTTGAACTGGTACATGGCGGCT |
| *RUBY_5GT*_qPCR_F | CTTATGGCACATCCGCCTACGT |
| *RUBY_5GT*_qPCR_R | TATGGTTCTCTGGGAAGCCTGG |
| *Rhg1-GmAAT*_qPCR_F | CGTGTAGAGTCCTTGAAGTACAGC |
| *Rhg1-GmAA*T_qPCR_R | ACCAGAGCTGTGATAGCCAACC |
| *GmEF1A*_qPCR_F | GACCTTCTTCGTTTCTCGCA |
| *GmEF1A*_qPCR_R | CGAACCTCTCAATCACACGC |
| **Cloning primers** | |
| *RUBY* cloning to pICH47802, F | gcagaagacaattgcgaattcaagcttggaggtcaacat |
| *RUBY* cloning to pICH47802, R | tgtgaagacaatgccagcgatctggattttagtactggat |
| *Rhg1-GmAAT* W370A, F | GCAATCCCAGATATTGCTTACTTCTTTCAGTTC |
| *Rhg1-GmAAT* W370A, R | GAACTGAAAGAAGTAAGCAATATCTGGGATTGC |
| *Rhg1-GmAAT* W370I,F | GCAATCCCAGATATTATTTACTTCTTTCAGTTC |
| *Rhg1-GmAAT* W370I,R | GAACTGAAAGAAGTAAATAATATCTGGGATTGC |
| *Rhg1-GmAAT* Y268L, F | GCTTTGTGCTGTGATCTTGCTTGCAATAGGCTTATTTG |
| *Rhg1-GmAAT* Y268L, R | CAAATAAGCCTATTGCAAGCAAGATCACAGCACAAAGC |
| *Rhg1-GmAAT* Y268A, F | GCTTTGTGCTGTGATCGCTCTTGCAATAGGCTTATTTG |
| *Rhg1-GmAAT* Y268A, R | CAAATAAGCCTATTGCAAGAGCGATCACAGCACAAAGC |
| *Rhg1-GmAAT* Y268F, F | GCTTTGTGCTGTGATCTTCCTTGCAATAGGCTTATTTG |
| *Rhg1-GmAAT* Y268F, R | CAAATAAGCCTATTGCAAGGAAGATCACAGCACAAAGC |
| *Rhg1-GmAAT* D286A, F | GGATTCAACCCAGTCAGCTATTCTCATCAATTTTG |
| *Rhg1-GmAAT* D286A, F | CAAAATTGATGAGAATAGCTGACTGGGTTGAATCC |
| *Rhg1-GmAAT* D286F, F | GGATTCAACCCAGTCAttcATTCTCATCAATTTTG |
| *Rhg1-GmAAT* D286F, R | CAAAATTGATGAGAATgaaTGACTGGGTTGAATCC |
| *Rhg1-GmAAT* D122A, F | CCTTATCATCATCGGAgctGTGCTATCTGGAAAGC |
| *Rhg1-GmAAT* D122A, F | GCTTTCCAGATAGCACagcTCCGATGATGATAAGG |
| *Rhg1-GmAAT* D122F, F | CCTTATCATCATCGGAttcGTGCTATCTGGAAAGC |
| *Rhg1-GmAAT* D122F, R | GCTTTCCAGATAGCACgaaTCCGATGATGATAAGG |
